# Marsupial single-cell transcriptomics provides an atlas of developmental heterochrony

**DOI:** 10.1101/2024.05.26.595746

**Authors:** Sergio Menchero, Christopher Barrington, Gregorio Alanis-Lobato, Wazeer Varsally, Kathy K. Niakan, James M. A. Turner

## Abstract

Single-cell transcriptomics has revealed conserved and divergent programmes of organogenesis in mammals, but existing studies have focused on eutherians. Marsupials exhibit short gestation and complete development externally, necessitating accelerated differentiation of anterior features required for locomotion and feeding. As such, they represent a unique outgroup with which to understand temporal shifts in development, known as heterochrony. Here, we generate the first single-cell transcriptomic atlas of gastrulation and early organogenesis in a marsupial, the opossum *Monodelphis domestica*. We find that anterior prioritisation is achieved by earlier initiation and shorter duration of transcriptional programmes relative to eutherians. The result is uncoupling of transcriptional and morphological progression, revealing unforeseen diversity in the order of developmental sequences in mammals. We uncover novel tissues for which heterochrony has not previously been documented. Our findings indicate that accelerated activation of transcriptional programmes facilitates the rapid growth needed for survival of the marsupial neonate.

## Introduction

Embryonic development entails the emergence of different cell types driven by the activation of specific genetic and cellular programmes. In vertebrates, the gene regulatory networks and morphogenetic changes that drive these transitions are broadly conserved ^1–3^. Nevertheless, in many species, there is a shift in the rate, timing, or order in which certain cell types or tissues arise. This change in timing relative to a related species is known as heterochrony ^4,5^. Developmental heterochrony has been proposed as an evolutionary force that results in phenotypic diversification, with examples across multiple phyla. For instance, heterochrony between the formation of the head and trunk has been linked to the evolution of feeding larvae in bilateria ^6^. An increased rate in the formation of somites results in a higher number of vertebrae typical of snakes ^7^. The delayed and protracted maturation of cortical synapsis is characteristic of the human brain ^8,9^. Despite its implications, the temporal diversification of tissue programmes throughout evolution is understudied.

Marsupial (metatherian) neonates exhibit short gestation compared to other mammals, and development is completed externally after birth. As a consequence, they undergo a marked anterior to posterior (A-P, rostral to caudal) gradient in development, with a particularly prominent growth of facial structures and forelimbs ^10^. This anterior-posterior heterochrony allows altricial newborns to crawl and receive nutrients from the mother. This phenomenon has been appreciated for decades at the morphological level but less so at the transcriptional level. Existing studies have focused on neural crest and limb formation, where early expression of selected genes has been reported ^11–14^. High resolution studies could identify whether anterior prioritisation is achieved only via early initiation or, in addition, by reduced duration of developmental processes. Furthermore, they could identify other tissues for which heterochrony has not been documented.

The expansion of single-cell technologies applied to cells from different tissues, embryonic time points and species has generated transcriptional atlases of development ^15–25^. Cross-species comparisons are particularly powerful at unveiling conserved signatures and specific evolutionary shifts. For instance, previous studies have identified conserved core pluripotency factors and divergent primordial germ cell expression programs in mammalian embryos ^18,21,26–28^.

To gain insight into developmental heterochrony, we performed single-cell RNA-sequencing (scRNA-seq) analysis of gastrulation and early organogenesis in grey short-tailed opossum (*Monodelphis domestica*) embryos, and compared this to equivalent eutherian datasets. Our analyses reveal the transcriptomic landscape of neural crest and forelimb specification, and identify additional tissues exhibiting early differentiation, such as the spinal cord and the anterior endoderm. Cross-species comparisons and RNA fluorescent *in situ* hybridization chain reaction (HCR RNA-FISH) experiments revealed early specification and rapid progression in the transcriptional cascade of specific tissues relative to underlying morphological landmarks, such as neural tube closure or limb bud outgrowth. Our findings implicate transcriptional control as an important mechanism in developmental heterochrony.

## Results

### Cross-species transcriptional equivalence

Gestation in the opossum lasts 14 days, with implantation around day 11-12. Using scRNA-seq, we previously showed that the first week of gestation culminates in the segregation of the three blastocyst lineages: epiblast, trophectoderm and primitive endoderm ^29^. At embryonic day (E) 11.5, the anterior structures of the opossum embryo, including head and forelimbs, are morphologically advanced ^30^ (**Figure S1A**). We therefore focused on intermediate time points (E8.5, E9.5, E10.0, and E10.5) when gastrulation and early organogenesis take place (**Figure 1A** and **1B**). Several embryos from the same litter were pooled by stage and processed for scRNA-seq using the droplet-based 10X Genomics Chromium platform. To gain spatial resolution, embryos at E10.0 and E10.5 were divided into two parts along the A-P axis. After quality control, 23,669 cells, were retained for analysis (**Figures S1B** and **S1C, Table S1 and S2**).

**Figure 1.**
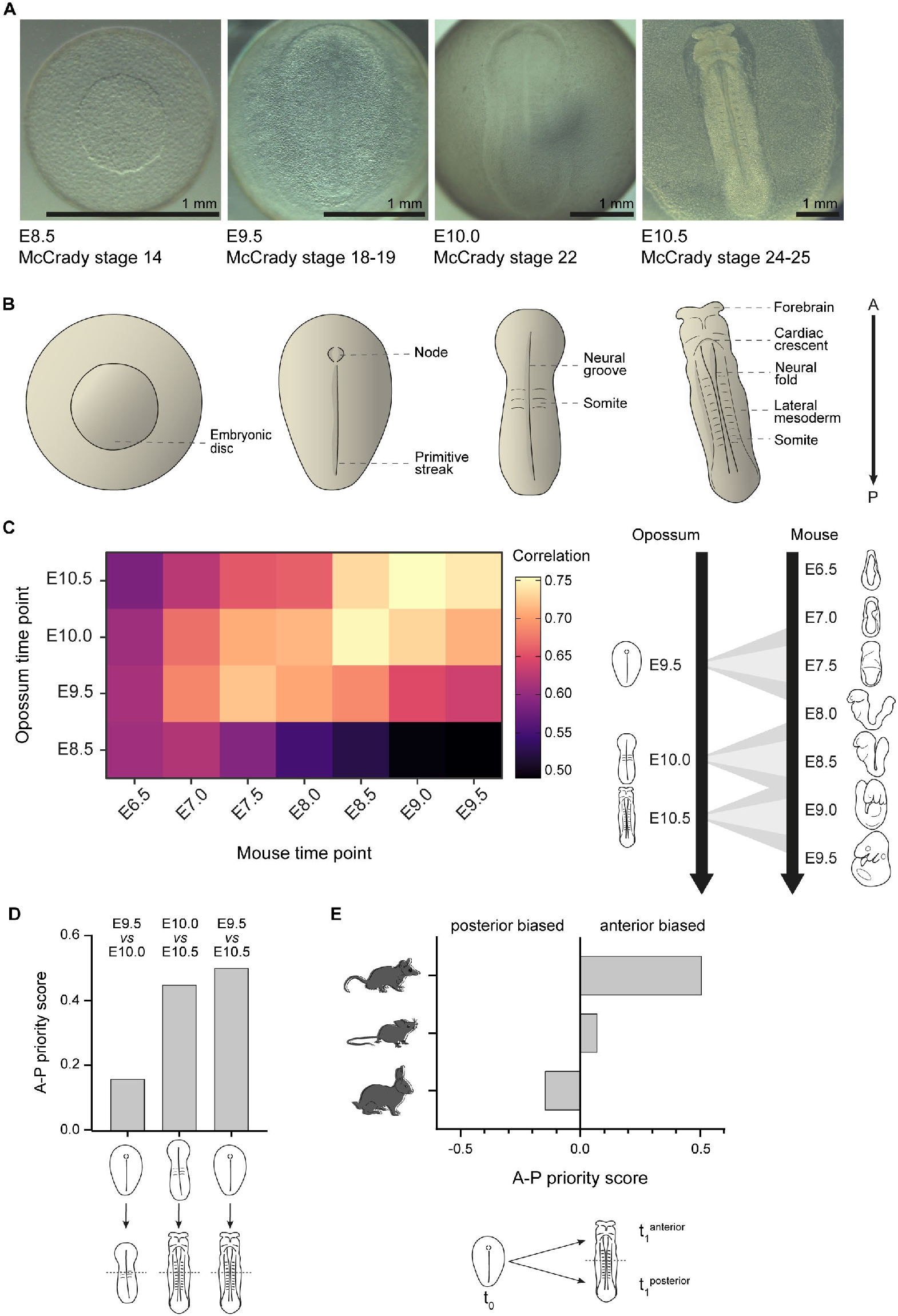
Cross-species transcriptional equivalence. (A) Stereo microscope images of opossum embryos at E8.5, E9.5, E10.0 and E10.5. Scale bars, 1 mm. (B) Schematic illustrations of opossum embryos with key features annotated. (C) Heatmap of Spearman’s rank correlation coefficients of gene expression per opossum (vertical) and mouse (horizontal) time point. The intersect of variable features across time points between both species is used to calculate the correlation. Schematic diagrams showing the transcriptional correspondences are summarised on the right panel. (D) A-P priority score for opossum (t0 = E9.5; t1 = E10.5), mouse (t0 = E7.5; t1 = E8.5, E9.0 and E9.5) and rabbit (t0 = E7; t1 = E9) embryos. (E) A-P priority score for opossum embryos using different starting and ending time points (t0 = E9.5, t1 = E10.0; t0 = E10.0, t1 = E10.5; and t0 = E9.5, t1 = E10.5).

Cross-species comparisons of transcriptional states in embryos can be useful to establish stage equivalences and assess developmental timing ^18,21,25,31^. In order to place our opossum dataset in context, we used available mouse datasets as a reference ^17,19^ and addressed developmental correspondence between species. After removing cell clusters associated with extraembryonic tissues (**Figure S1D** and **S1E, Table S3**), we pseudobulked all cells from each time point. We defined a set of one-to-one orthologues and identified variable features across the timecourse common to both species to focus on genes informative for developmental progression. Opossum E8.5 embryos had a low correlation with all mouse time points (**Figure 1C**), as they correspond to pre-primitive streak stages. Opossum E9.5 embryos were most similar to mouse E7.5 embryos, when both species are at the late primitive streak stage (**Figure 1C**). Remarkably, opossum E10.0 and E10.5 embryos exhibited a transcriptomic profile that in mice is found at a more advanced morphological state. At E10.0, the opossum has 1-2 somites, neurulation has not started, and there is no visible heart, yet the transcriptome resembled that of an E8.5 mouse, in which there are 6-10 somites, the neural tube is starting to close, and the heart is beating (**Figure 1C**). At E10.5, the opossum is relatively flat, has 10-12 somites, the limb buds are not evident, the neural tube is starting to close, and the heart is beating (but not fused in the midline). However, the transcriptome resembled that of an E9.0 mouse, when there are 16-19 somites, the forelimb buds are present, the neural tube is closed, the lens vesicle is forming, the gut tube is closed, and the heart is looping (**Figure 1C**). We observed a similar phenomenon when comparing opossum to rabbit ^21^ and macaque embryos ^25^: opossum E10.5 embryos corresponded transcriptionally to rabbit E9 embryos (**Figure S2A**) and to macaque E26-E29 embryos (**Figure S2B**). At these stages in rabbit and macaque embryos, the neural tube is closed, the heart is prominent, the optic vesicle is evident and the forelimb bud is emerging ^32,33^. To deconvolve the origin of this similarity, we compared our opossum anterior and posterior datasets individually to these eutherian species. We observed that the correlation coefficient was higher for the anterior than the posterior opossum dataset when compared to morphologically advanced eutherian stages (**Figure S2C**).

To delineate when during opossum development anterior structures become prioritised over posterior ones, we calculated an “A-P priority” score, in which we calculated the number of differentially expressed genes between either the anterior or posterior datasets and an earlier timepoint (**Figure S2D**). A-P prioritisation was clearly evident between E10.0 and E10.5: the A-P score was + 0.16 in the E9.5 - E10.0 comparison, but increased to + 0.45 in the E10.0 to E10.5 comparison. Consequently, the A-P score was also higher in the E9.5 to E10.5 comparison **(Figure 1D**). These results indicate a greater change in gene expression in the anterior than the posterior embryo region from the onset of organogenesis. To contextualize this with eutherians, we used mouse ^17^ and rabbit ^21^ anterior and posterior datasets. In the mouse, we compared E7.5 with E8.5, E9.0 and E9.5 anterior and posterior datasets. The A-P priority score indicated a more subtle bias towards the anterior in all cases (+ 0.07 on average) (**Figure 1E, Figure S2E**). In the case of the rabbit, we compared E7 with E9. The A-P priority score in this case was negative (- 0.15) (**Figure 1E**), suggesting a bias towards posterior tissues. These analyses demonstrate uncoupling between the transcriptional and morphological progression of opossum embryos, mainly due to the rapid transcriptional differentiation of the anterior part of the embryo at the onset of organogenesis.

### Conserved transcriptional progression at gastrulation

Next, we used uniform manifold approximation and projection (UMAP) to visualise clusters of cell types within our single-cell data (**Figures 2A-2D**). To aid assignment of cell types to each cell cluster, we calculated a gene module score based on the combinatorial expression of conserved key marker genes associated with those cell types/states (**Table S3** and **S4, Figures S3, S4** and **S5**). At E8.5, we found the same clusters that are present at E7.5 ^29^: the embryonic epiblast, trophectoderm and primitive endoderm (**Figure 2A, Figures S1D, S1E** and **S3A**). Pluripotency-associated markers *POU5F1, POU5F3* and *NANOG* were detected in most epiblast cells. Expression of genes that in the chicken and mouse precede primitive streak formation, e.g., *FGF8, PRICKLE1, CNOT1*, and *TBXT*, was detected in a few epiblast cells (**Figure 2E**) ^34–37^. Immunofluorescence and fluorescent RNA *in situ* hybridization chain reaction (HCR IF + RNA-FISH) revealed that POU5F3 was expressed throughout the epiblast, while *FGF8* expression was polarised to one side of the embryonic disc (**Figure 2F**). This indicates that symmetry is broken at the molecular level by E8.5, consistent with a previous study ^38^.

**Figure 2.**
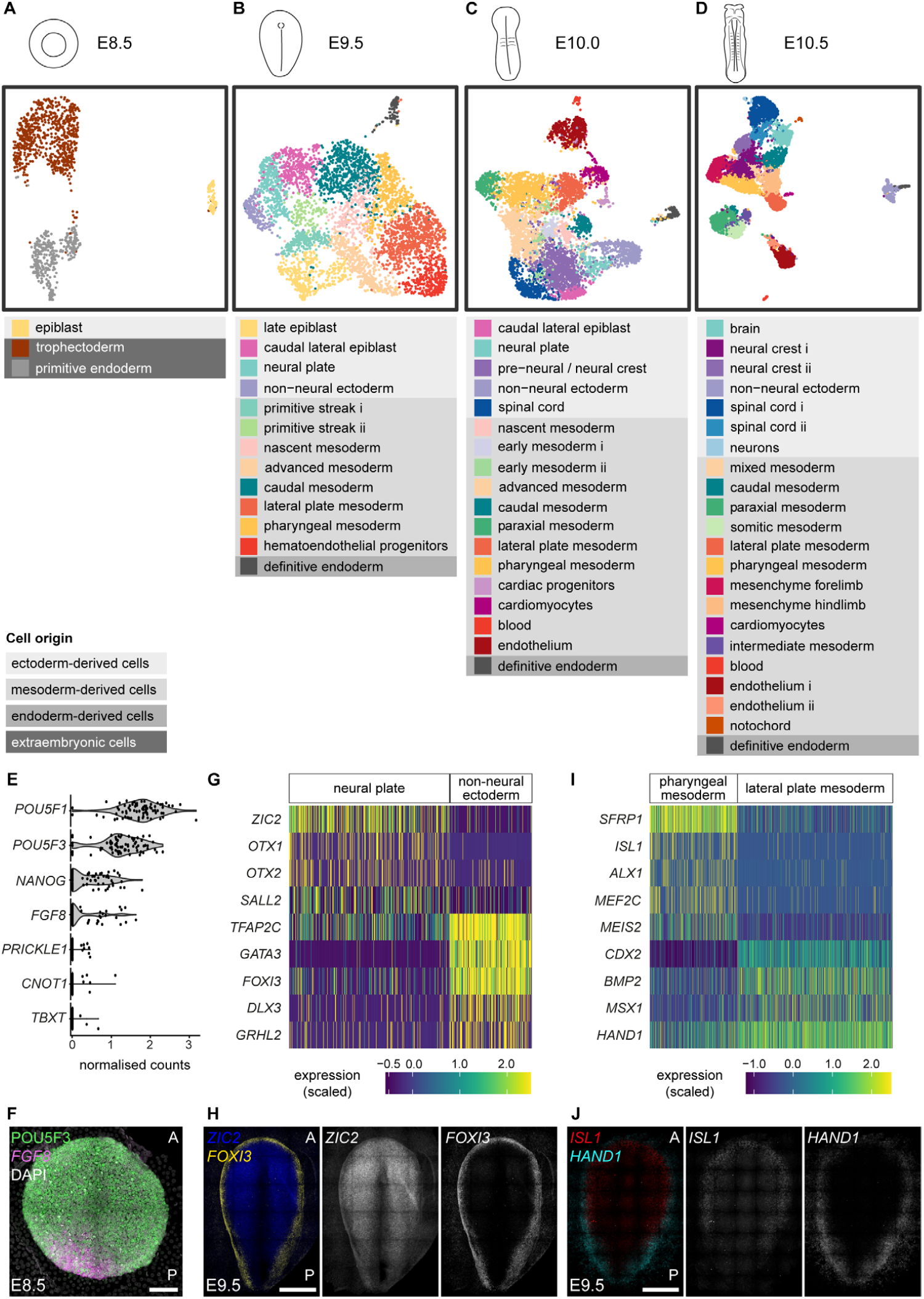
Opossum embryo single-cell clustering and annotation. (A-D) UMAP of opossum scRNA-seq data from E8.5, E9.5, E10.0 and E10.5 embryos. Clusters were annotated using module scores. Module scores are based on the combinatorial expression of markers reported in Table S3. (E) Violin plots for the expression of pluripotency (*POU5F1, POU5F3, NANOG*) and pre-primitive streak markers (*FGF8, PRICKLE1, CNOT1, TBXT*) in the epiblast of E8.5 embryos. (F) Expression of POU5F3 (HCR immunofluorescence) and *FGF8* (HCR RNA-FISH) in the epiblast of an E8.5 embryo. Nuclei are stained with DAPI. Image is a maximal projection. A = anterior; P = posterior. Scale bar, 100 µm. (G) Heatmap of genes enriched in neural plate and non-neural ectoderm at E9.5. (H) Expression of *ZIC2* (neural plate) and *FOXI3* (non-neural ectoderm) by HCR RNA-FISH in an E9.5 embryo. Images are maximal projections. A = anterior; P = posterior. Scale bar, 500 µm. (I) Heatmap of genes enriched in pharyngeal mesoderm and lateral plate mesoderm at E9.5. (J) Expression of ISL1 (pharyngeal mesoderm) and HAND1 (lateral plate mesoderm) by HCR RNA-FISH in an E9.5 embryo. Images are maximal projections. A = anterior; P = posterior. Scale bar, 500 µm.

Transcriptome comparison of mouse and opossum embryos highlighted that the late primitive streak stage (mouse E7.5 and opossum E9.5) was similar between both species (**Figure 1C**). In the mouse, different subtypes of ectoderm and mesoderm are segregated at this point (E7.5) ^19^. In the opossum, our clustering approach also identified distinct ectoderm and mesoderm subtypes at E9.5 (**Figure 2B**). Neural and non-neural ectoderm were segregated, and the expression of key genes such as *ZIC2* and *OTX2* (neural plate) or *TFAP2C, GATA3* and *FOXI3* (non-neural ectoderm) was enriched in their corresponding clusters (**Figure 2G**) ^39–43^. Similarly, the most anterior part of the opossum lateral plate mesoderm, the pharyngeal mesoderm, had acquired a specific transcriptional signature (**Figure 2B**). This included the expression of cardiac progenitor markers such as *MEF2C, ISL1* and *MEIS2* ^44–46^, that distinguished it from the most posterior lateral plate mesoderm (**Figure 2I**). Using HCR RNA-FISH in whole-mount embryos, we verified that these ectoderm and mesoderm subtypes were also spatially segregated as expected (**Figures 2H** and **2J**). Thus, mouse E7.5 and opossum E9.5 embryos exhibit broadly similar cell types.

### Advanced transcriptional progression at organogenesis

While opossum embryos exhibited a similar profile to the mouse until the late primitive streak stage, cross-species comparisons revealed an advanced transcriptional progression at organogenesis (**Figure 1C**). At E10.0, we found transcriptional signatures corresponding to some advanced cell types, including paraxial mesoderm, cardiomyocytes, spinal cord, and neural crest (**Figure 2C**). It was remarkable to find cells with a neural crest signature given the flat morphology of the opossum embryo at this point. *Bona fide* neural crest cells are only found in the mouse when the body plan is more advanced, around E8.25-E8.5, ^19^.

By E10.5, clusters that defined mature cell types became a prominent feature of the dataset. Forelimb and hindlimb domains were segregated, the neural crest and spinal cord clusters were subdivided, and even a cluster of neurons was identified (**Figure 2D**), something that only happens in the mouse when the neural tube has been fully closed around E9.5 ^17,20^.

These findings support the transcriptional equivalence determined between mouse and opossum embryos (**Figure 1C**), and confirm accelerated developmental progression at the transcriptome level at the onset of organogenesis in the opossum. To better understand this rapid progression, we focused on tissues whose early differentiation is important for the survival of the marsupial neonate.

### Early specification and migration of the neural crest

The neural crest is a migratory multipotent cell population that is specified in the ectoderm, at the border between the neural tube and non-neural ectoderm, and gives rise to a wide variety of cell types including bone, cartilage, neurons, glia, and melanocytes ^47^. In vertebrates, neural crest precursors are induced during gastrulation ^48,49^ but bona fide neural crest cells, characterised by the expression of *SOX9* and *SNAI1*/*2* among others, are only found once neurulation begins ^50^. Subsequently, they upregulate *SOX10*, delaminate, and migrate to their definitive domains. In mammals, delamination of the most-anterior neural crest cells, the cranial neural crest, occurs prior to neural tube closure while delamination in the trunk and posterior part of the embryo happens after neural tube closure ^51^. Eventually, around E9.5 in the mouse, cranial neural crest cells populate the frontonasal prominence and pharyngeal arches, where they acquire an ectomesenchyme fate and contribute to the craniofacial skeleton ^52^.

Given that marsupials exhibit rapid development of craniofacial structures, we characterised when the neural crest is specified and when it differentiates. We identified subpopulations of anterior neural crest using conserved markers of the gene regulatory network ^47,53^ and validated our findings using HCR RNA-FISH. Our analysis revealed interesting similarities and differences from eutherians. We identified four neural crest subpopulations and interrogated the expression of key genes in each (**Figure 3A, Figure S6A**). The first, formed mostly at E10.0, was enriched with neural crest specification and delamination markers, including *TFAP2C, SOX9* and *SNAI2* (**Figure 3A, Figure S6B**). While this finding suggests that the neural crest is specified at E10.0, *SOX9* was already expressed in the anterior neural plate at E9.5 (**Figure 3B**). This is earlier than observed in mouse and chicken, in which *SOX9* is first expressed around the time of somitogenesis ^14^, and indicates that neural crest specification occurs particularly early in the opossum. At E10.0, *SOX9* was confined to the border of the neural plate, consistent with a previous study ^14^. Although *SOX9* was more abundant in the most anterior part, the domain extended to about 2/3 of the embryo length (**Figure 3C**). A second cluster was enriched in *SOX10* and *NGFR*, genes associated with cell migration (**Figure 3A**). HCR RNA-FISH confirmed that *SOX10* is expressed from E10.0, but did not extend as posteriorly as *SOX9* (**Figure 3C**) indicating that, as in eutherians, neural crest differentiation progresses in an anterior to posterior manner. At E10.5, anterior neural crest cells had migrated and populated the pharyngeal arches (**Figure 3D**). However, in contrast to eutherians, trunk neural crest cells had initiated delamination and migration prior to neural tube closure (**Figures 3D** and **3E**).

**Figure 3.**
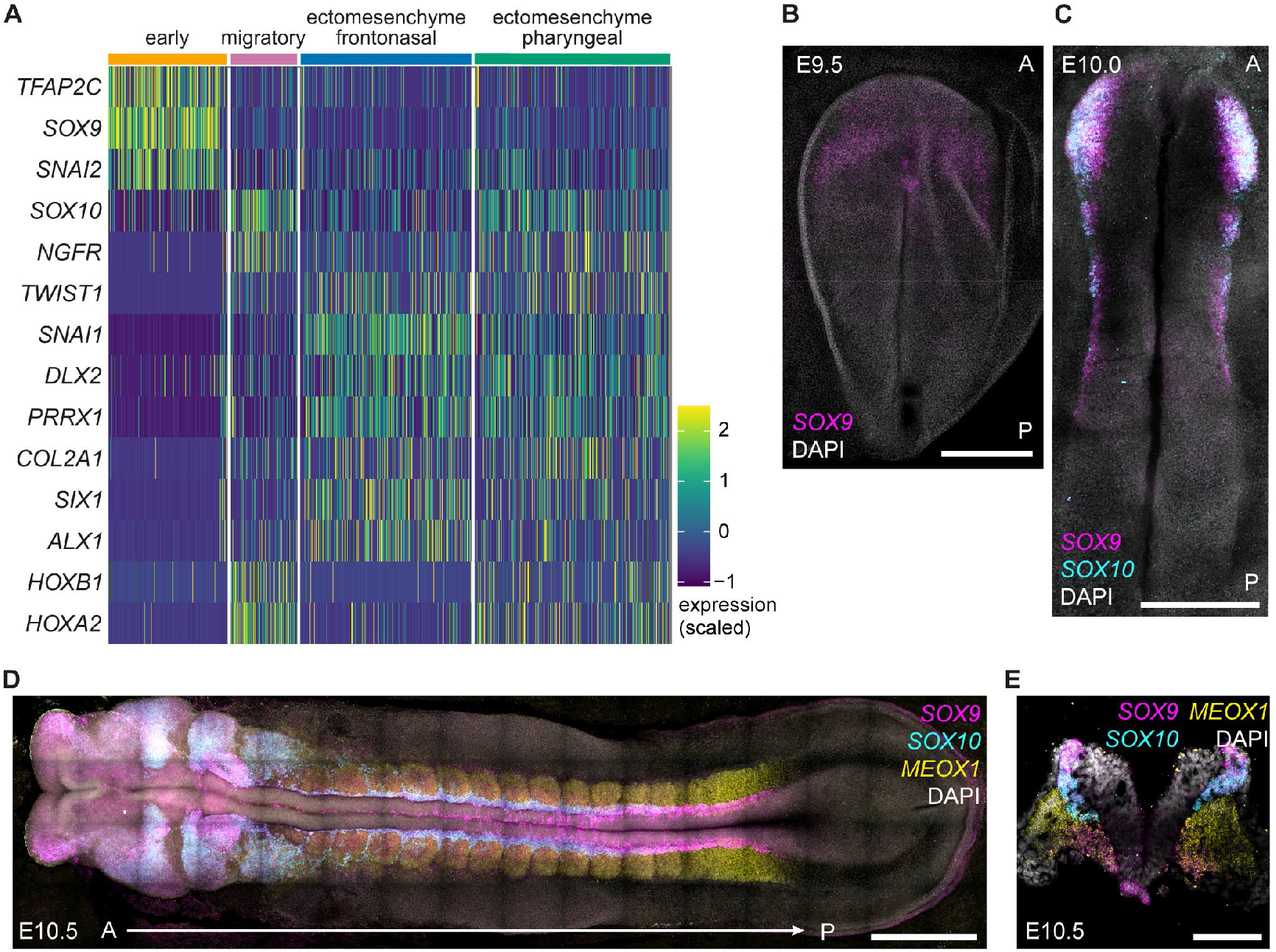
The opossum neural crest. (A) Heatmap of genes enriched in different clusters of anterior neural crest cells from opossum E10.0 and E10.5 embryos. (B) Expression of *SOX9* by HCR RNA-FISH in an E9.5 embryo. Nuclei are stained with DAPI. Image is a maximal projection. A = anterior; P = posterior. Scale bar, 500 µm. (C) Expression of *SOX9* and *SOX10* by HCR RNA-FISH in an E10.0 embryo. Nuclei are stained with DAPI. Image is a maximal projection. A = anterior; P = posterior. Scale bar, 500 µm. (D) Expression of *SOX9, SOX10* and *MEOX1* by HCR RNA-FISH in an E10.5 embryo. Nuclei are stained with DAPI. Image is a maximal projection. A = anterior; P = posterior. Scale bar, 500 µm. (D) Transversal section showing expression of *SOX9, SOX10* and *MEOX1* by HCR RNA-FISH in an E10.5 embryo. Nuclei are stained with DAPI. Image is a maximal projection. Scale bar, 100 µm.

The two remaining neural crest clusters exhibited an enrichment of transcripts linked to an ectomesenchyme fate, such as *TWIST1, SNAI1* and *PRRX1* (**Figure 3A**). One cluster was enriched in the frontonasal marker *ALX1*, while the other expressed the pharyngeal arch markers *HOXB1* and *HOXA2* (**Figure 3A**). This result indicated that frontonasal and pharyngeal subtypes are present 12 hours after the specification of neural crest cells. This contrasts with the mouse, in which the same process takes around 24 hours ^53^. We conclude that the prominent differentiation of craniofacial features in marsupials is facilitated by both early initiation and subsequent rapid differentiation of the neural crest.

### Dorsal-ventral patterning and first neurons in the spinal cord

The spinal cord forms the posterior region of the central nervous system and is the source of neurons that process sensory information and coordinate motor activity. During development, the spinal cord undergoes dorsal-ventral patterning, induced by opposing morphogen gradients originating from the adjacent non-neural ectoderm and the notochord ^54–57^. These gradients induce expression of transcription factors including *MSX1*/*2, PAX3*, and *NKX6-1* within restricted domains along the dorsal-ventral axis, leading to specification of distinct neural progenitors ^58^. These progenitors then differentiate into specialised post-mitotic neurons. In mouse, dorsal-ventral patterning of the spinal cord begins before neural tube closure. However, the differentiation of neural progenitors to neurons only occurs after neural tube closure, around E9.0-E9.5 in the mouse.

We investigated whether dorsal-ventral patterning of the spinal cord was evident in our opossum dataset and found similarities and differences from eutherians. Analysis of the integrated per time point datasets identified one spinal cord cluster at E10, and two spinal cord and one neuron clusters at E10.5 (**Figures 2C** and **2D**). We extracted these cells and re-clustered them agnostic to gestational age, identifying two spinal cord clusters and one neuron cluster (**Figures S7A** and **S7B**). Interestingly, the two spinal cord clusters exhibited differences in *HOX* gene expression, indicating that we had identified anterior and posterior spinal cord populations ^59^. The presumptive anterior cluster was enriched for *HOXB4* expression, which is characteristic of the anterior cervical region, while the presumptive posterior cluster was enriched for *HOXC6, HOXA7, HOXB7*, and *HOXB9*, which mark the posterior cervical and thoracic regions of the spinal cord (**Figure 4A**) ^12,60^.

**Figure 4.**
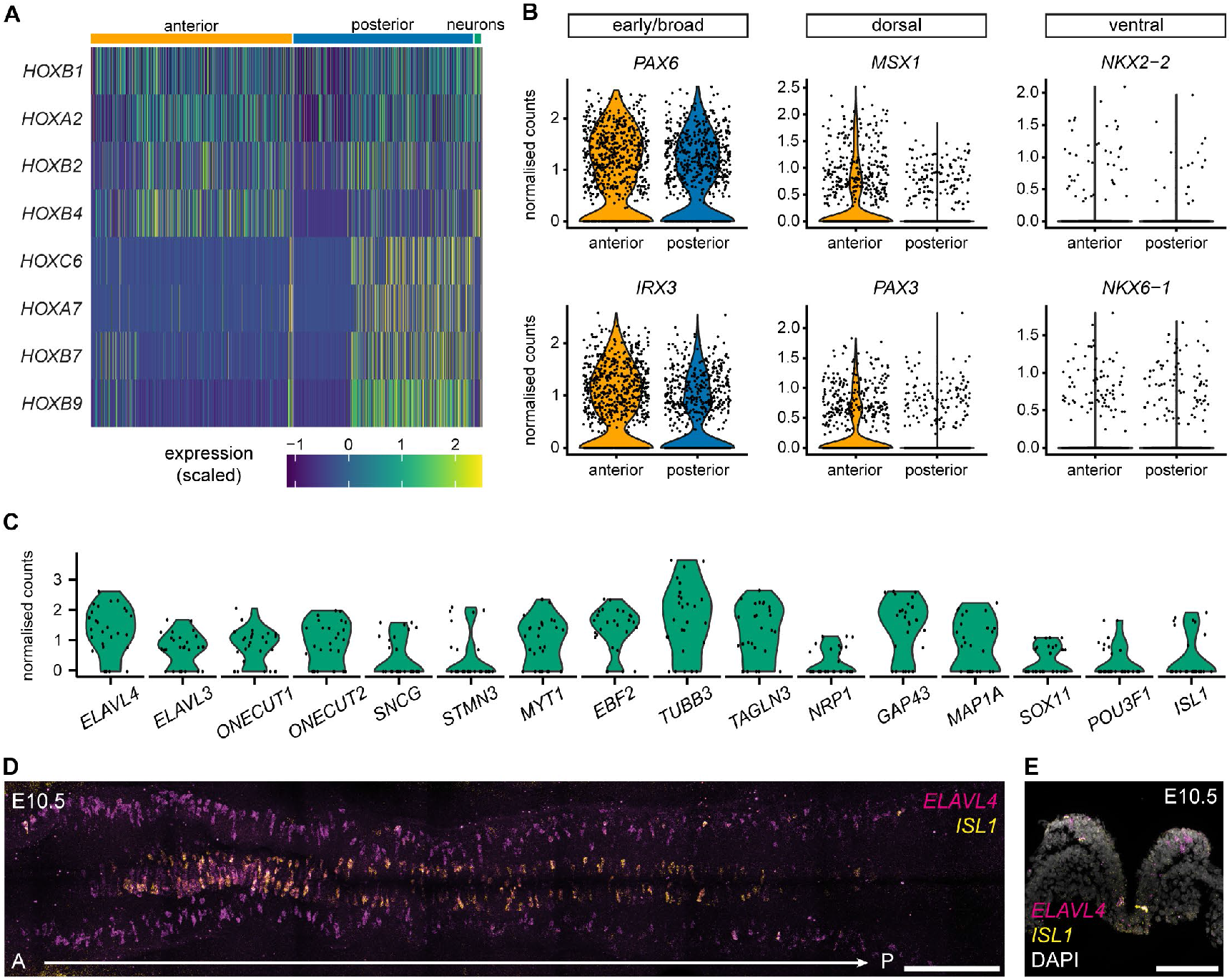
Patterning of the opossum spinal cord. (A) Heatmap of *HOX* gene expression in spinal cord and neuron clusters from opossum E10.0 and E10.5 embryos. (B) Violin plots for the expression of early/broad (*PAX6, IRX3*), dorsal (*MSX1, PAX3*) and ventral (*NKX2-2, NKX6-1*) spinal cord markers in anterior and posterior populations. (C) Violin plots for the expression of neuron-associated markers in opossum neurons at E10.5. (D) Expression of neuronal markers *ELAVL4* and *ISL1* by HCR RNA-FISH in the spinal cord of an E10.5 embryo. Image is a maximal projection. A = anterior; P = posterior. Cells in the midline represent ventral positions while cells on the lateral correspond to dorsal regions. Scale bar, 200 µm. (E) Transversal section showing expression of *ELAVL4* and *ISL1* by HCR RNA-FISH in the spinal cord of an E10.5 embryo. Nuclei are stained with DAPI. Image is a maximal projection. Scale bar, 100 µm.

We then examined whether genes that define distinct domains in the dorsal-ventral axis were differentially expressed in the anterior and posterior spinal cord clusters ^61,62^. Both clusters expressed dorsal (*MSX1* and *PAX3*) and ventral (*NKX2-2* and *NKX6-1*) markers (**Figure S7C**). However, the proportion of cells expressing dorsal markers was higher in the anterior population, suggesting that dorsal-ventral patterning is more established in the anterior versus the posterior region of the spinal cord (**Figure 4B** and **Figure S7D**). We conclude that, similarly to eutherians, cells respond to the opposing morphogen gradient prior to neural tube closure and the resulting patterning advances in an anterior-to-posterior manner.

We then investigated the transcriptional signature of the neuron cluster. We observed expression of known pan-neuronal markers, including *ELAVL3*/*4, ONECUT1*/*2, TUBB3, MYT1* and *GAP43* (**Figure 4C**) ^61,62^. These neurons lacked neural crest-derived neuron markers, confirming that they derive from the central nervous system (**Figure S7E**). In addition, *ISL1*, a marker of motor neurons and a dorsal interneuron population, was expressed in this cluster. HCR RNA-FISH confirmed that *ISL1*-positive cells were particularly enriched ventrally, in the presumptive motor neuron domain (**Figure 4D** and **4E**). This finding was surprising because, in mouse and chicken, *ISL1* expression is detected after the spinal cord is closed and the number of neural progenitors is higher ^63,64^. Furthermore, we observed a higher density of neurons in the anterior part of the spinal cord, consistent with most neurons expressing *HOXB4* (**Figure 4A** and **4D**). These findings reveal that in the opossum spinal cord, neural progenitor differentiation is accelerated, giving rise to post-mitotic neurons in less than 12 hours. This differs from the mouse, in which the same process takes at least 24 hours ^65^.

### Early patterning precedes outgrowth in the forelimb domain

Limb formation begins with the specification of the limb fields in the lateral plate mesoderm at specific positions along the A-P axis. The transcription factor *TBX5* determines the forelimb domain, while *PITX1* and its downstream target *TBX4* determine hindlimb identity ^66–70^. These transcription factors activate *FGF10* in the mesoderm, which then activates *FGF8* in the apical ectodermal ridge to create a positive FGF-signalling feedback loop that promotes limb outgrowth ^71–73^. Expression of *HOX* genes in the lateral plate mesoderm determines the position of the forelimb, the interlimb flank and the hindlimb ^74^. In most mammals, formation of the forelimb and hindlimb is asynchronous, with the hindlimb relatively delayed ^75–77^. For instance, in the mouse, the forelimb bud is first evident at E9.0-E9.5, while the hindlimb bud is not until E10.0 ^78^.

Forelimb-hindlimb heterochrony is extreme in the case of marsupials, which quickly develop their forelimbs to crawl to the teats of the mother after birth ^10^. Based on marker analysis, this heterochrony has been attributed to differences in the timing of limb bud outgrowth, rather than to a delay in hindlimb specification ^12^. Consistent with this model, at E10.5, before any limb bud is obvious, we identified signatures that in other vertebrates typify forelimb and hindlimb identity, including expression of *TBX5* in the forelimb and *PITX1* and *ISL1* in the hindlimb. Nevertheless, transcriptional comparisons revealed that the forelimb and hindlimb were developmentally asynchronous (**Figure 5A**). The forelimb field expressed markers associated with later stages of limb development, including *FGF10, INKA1, MECOM* (also known as *EVI1*) and *IRX3* ^72,79–82^. Conversely, the hindlimb retained expression of the lateral plate mesoderm markers *HAND1* and *GATA6*, and had not yet upregulated *TBX4*. We used HCR RNA-FISH for *TBX5* and *PITX1* to locate the opossum forelimb and hindlimb domains, respectively. At E10.5, *TBX5* was detected in the cardiac crescent and presumptive forelimb domain, while *PITX1* was expressed in the posterior lateral plate mesoderm marking the presumptive hindlimb domain (**Figure 5B**). In the mouse, *TBX5* is initially expressed in a broader domain than the future forelimb-forming region ^83^. However, as previously reported, the opossum *TBX5* domain was more expanded ^12^, extending posteriorly to the level of the presomitic mesoderm, and abutting the *PITX1* domain. As a consequence, and unlike in chicken and mouse ^67,69^, no interlimb region was observed at E10.5. Therefore, forelimb development is characterised by both heterochrony and heterotopy, i.e. a shift in the spatial patterning of a developmental feature.

**Figure 5.**
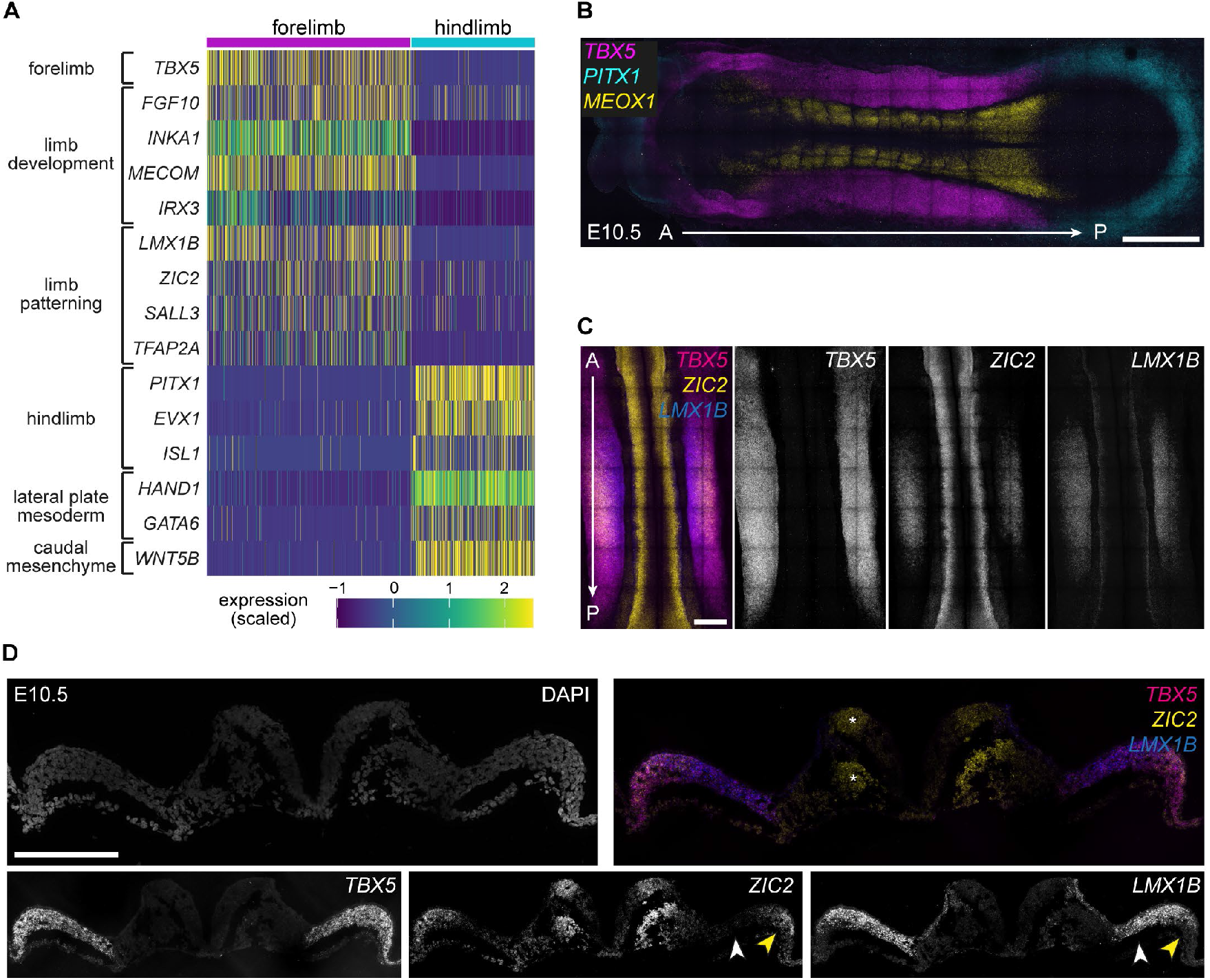
Limb patterning in the opossum embryo. (A) Heatmap of genes enriched in the forelimb and hindlimb domains at E10.5. (B) Expression of *TBX5* (heart and forelimb), *PITX1* (hindlimb) and *MEOX1* (somites) by HCR RNA-FISH in an E10.5 embryo. Image is a maximal projection. A = anterior; P = posterior. Scale bar, 500 µm. (C) Expression of *TBX5, ZIC2* and *FGF8* and in the forelimb of an E10.5 embryo. *ZIC2* marks the distal domain while *FGF8* is expressed in the limb ectoderm. Image is a maximal projection. A = anterior; P = posterior. Scale bar, 200 µm. (D) Transversal section showing expression of *TBX5* (broad), *LMX1B* (proximal, white arrowhead) and *ZIC2* (distal, yellow arrowhead) in the opossum E10.5 forelimb by HCR RNA-FISH. *ZIC2* is also expressed in dorsal spinal cord and somites (*). Nuclei are stained with DAPI. Images are maximal projections. Scale bar, 200 µm.

Morphological differences between the opossum forelimb and hindlimb become obvious at E11.5 ^30^ (**Figure S1A**). We addressed whether patterning and differentiation at the transcriptional level precede morphological changes, as observed in the spinal cord. In mouse and chicken, transcription factors associated with dorsal-ventral or proximal-distal patterning (e.g., *LMX1B, ZIC2, SALL3* and *TFAP2A* ^84–88^) are expressed after the limb bud is formed. Remarkably, in the opossum forelimb, these markers were detected at E10.5, prior to limb bud elongation (**Figure 5A**). Using HCR RNA-FISH we visualised local patterning in the forelimb domain at this stage. First, expression of *LMX1B* and *ZIC2* was restricted to the medial part of the *TBX5* domain, presumably anticipating the forelimb-forming territory (**Figure 5C**). This region coincided spatially with the expression of *FGF8* in the nascent apical ectodermal ridge (**Figure S8A**). In other vertebrates, restriction of *TBX5* to the definitive forelimb domain precedes expression of *LMX1B* and *ZIC2*. The regionalisation of these transcripts before *TBX5* in the opossum indicates that different factors are in place to demarcate the limb-forming territory. Second, we observed a proximal-distal patterning, with *LMX1B* enriched in the proximal area while *ZIC2* was segregated to the distal part of the forelimb (**Figure 5D**). These results indicate not only that the forelimb and hindlimb domains are specified early, but also that the forelimb undergoes accelerated patterning before limb bud outgrowth.

### Preferential patterning of anterior vs posterior endoderm

Finally, we investigated whether our transcriptome dataset would identify heterochrony in other, understudied tissues. Given that at birth marsupials require a respiratory and digestive system, we focused on endoderm differentiation. During gastrulation, definitive endoderm cells intercalate with the underlying extraembryonic endoderm cells to form a layer of cells in the most ventral part of the embryo ^89,90^. This sheet of cells then folds to give rise to an internal primitive gut tube that becomes regionalised along the A-P axis into foregut, midgut, and hindgut domains. These broad domains become gradually more specialised in subdomains, and organ buds emerge, giving rise to a variety of tissues including the thyroid, lung, liver, pancreas, and colon.

We investigated whether the definitive endoderm undergoes early patterning and differentiation. Using eutherian markers of different gut domains along the A-P axis ^91,92^, we generated module scores to assess the identity of the opossum definitive endoderm cells at E10.5 (**Table S5**). Most cells had a foregut or midgut identity but very few had a hindgut signature (**Figure 6A, Figure S9A**). We observed the same bias in extraembryonic-endoderm cells (**Figure S9B**), which are known to contribute to the gut endoderm ^92^.

**Figure 6.**
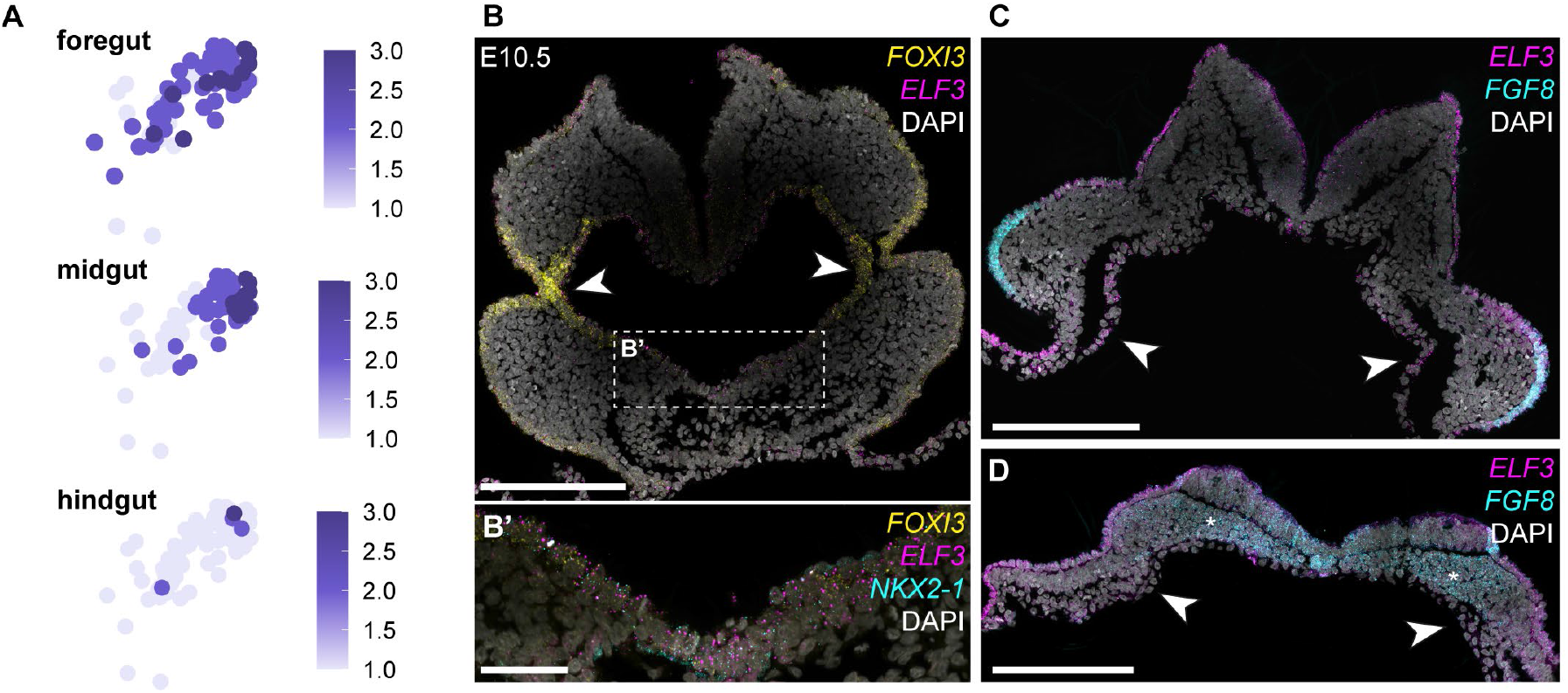
Definitive endoderm differentiation in the opossum. (A) Feature plots of E10.5 definitive endoderm cells showing enrichment of module scores for foregut, midgut, and hindgut signatures. Module scores are based on the combinatorial expression of markers reported in Table S5. (B) Expression of *FOXI3* and *ELF3* by HCR RNA-FISH in the anterior part of an E10.5 opossum embryo. Image is a maximal projection of a transversal section. Arrowheads indicate endoderm cells. FOXI3 also labels non-neural ectoderm cells. Scale bar, 200 µm. (B’) Expression of *FOXI3, ELF3* and *NKX2-1* in the ventral endoderm cells. Scale bar, 50 µm. (C) Expression of *ELF3* and *FGF8* by HCR RNA-FISH in the trunk of an E10.5 opossum embryo. Image is a maximal projection of a transversal section. *ELF3* is expressed in endoderm cells (arrowheads) and non-neural ectoderm cells. *FGF8* is expressed in limb ectoderm. Scale bar, 200 µm. (D) Expression of *ELF3* and *FGF8* by HCR RNA-FISH in the posterior part of an E10.5 opossum embryo. Image is a maximal projection of a transversal section. *ELF3* is expressed in non-neural ectoderm cells. FGF8 is expressed in presomitic mesoderm (*) but absent in endoderm (arrowheads). Scale bar, 200 µm.

Using HCR RNA-FISH we located the three gut domains in E10.5 embryos. We selected *FOXI3, ELF3*, and *FGF8* as markers of foregut, midgut, and hindgut respectively. *FOXI3* was expressed in the lateral region of the pharyngeal endoderm (**Figure 6B**, arrowheads) while *ELF3* was particularly enriched in the endoderm at the level of the forelimbs (**Figure 6C**, arrowheads). However, very few posterior endoderm cells expressed *FGF8*, consistent with hindgut formation being delayed relative to foregut and midgut (**Figure 6D**, arrowheads). The asynchronicity in the gut specification along its A-P axis was further emphasised by the expression of the lung and thyroid progenitor *NKX2-1* on the ventral side of the foregut (**Figure 6B’, Figure S9C**). This was remarkable given that, in the mouse, *NKX2-1* expression begins once the gut tube is fully closed, around E9.0 ^93^, while the opossum gut tube is only closed in the most anterior region at E10.5. Transcriptional analysis of the E10.5 endoderm revealed two clusters with differential A-P signatures (**Figure S9D**). The first was enriched in markers associated with thymus, thyroid and lung progenitors including *HMGB2, CRABP2* and *ATF3*, while the second was enriched in pancreas and small intestine progenitors such as *CPM, VAT1* and *DAB2* ^92^. These analyses reveal differences in the initiation timing of endoderm domains. While the foregut and midgut domains have acquired their specific transcriptional signature, the posterior gut is less differentiated.

## Discussion

The short gestation of marsupials has favoured the evolution of mechanisms prioritising the differentiation of tissues relevant for the immediate survival of neonates. In this work, we used the opossum as a model to provide insight into developmental heterochrony and the onset of tissue-specific prioritisation. Our findings reveal that this bias begins during early organogenesis, with rapid progression of tissues associated with craniofacial structures, locomotor activity, and respiratory and digestive systems. We also identify novel tissues for which heterochrony has not been previously documented.

The asynchronous development of anterior and posterior structures in marsupials has been appreciated for decades. Initially, the focus of this uncoupling was on the slow differentiation, or “dormancy”, of posterior structures given delayed hindlimb outgrowth ^12^ and the steep decrease of the segmentation clock to form posterior somites ^94^. However, the early migration of the cranial neural crest ^95^ and the early specification of the forelimb field ^11,12^ also pointed to acceleration of anterior structures. Our transcriptomic data support accelerated progression of anterior features as a key factor driving anterior bias. They also show that is achieved by both early initiation and reduced duration of developmental programs. The net effect is uncoupling of transcriptional and morphological progression, providing an interesting contrast to observations in eutherians. For instance, birth of post-mitotic neurons in the spinal cord can precede neural tube closure. Similarly, proximal-distal patterning of the limb field does not require limb outgrowth.

Species differences in the time of initiation of developmental programmes have been attributed to *cis*-regulatory elements. For instance, the early expression of *SOX9* in the opossum is driven by a marsupial-specific sequence in a *SOX9* enhancer ^96^. Our finding that transcriptional programmes are accelerated in tissues from all three germ layers suggests that broader mechanisms may also be at play. Differences in the pace of development between species has been linked to fundamental cellular functions, including the rate of protein degradation and metabolism ^97–101^. Whether these processes determine different developmental rates within the same organism is unclear.

Our work highlights how cross-species comparative transcriptomics can provide insight not only into the conservation of cell lineages but also into temporal shifts like heterochrony. Our findings show that marsupials deploy conserved developmental programmes but through a different temporal code. The datasets reported here provide a key resource for broader evolutionary analyses to delineate mammalian development.

## Methods

### Opossum maintenance

Opossums (*Monodelphis domestica*) were maintained following the United Kingdom Animal Scientific Procedures Act 1986, and according to ethical guidelines at the Francis Crick Institute. Protocols were approved by local ethical review and the UK Home Office (Project Licence PP1234304). Opossums were housed in individual double-decker cages (GR1800, Tecniplast), with constant access to water and food, on a 14/10 h light/dark cycle. During nonbreeding periods, males and females were maintained in different rooms at 24-28°C and 55-75% humidity. For breeding, male and female opossums were kept next to each other in their individual cages for two days, and then swapped into each other’s cages for two days. Subsequently, they were put together into the same cage and breeding was monitored by CCTV, which allowed accurate estimates of developmental stages.

### Opossum embryo recovery and single-cell dissociation

Embryos were recovered from the uterus of female opossums in PBS and kept on ice during processing. They were transferred to plates with HBSS (Life Technologies, 14175053) and 5% FBS for removal of the shell coat and most of the extraembryonic tissues (except for E8.5 embryos where extraembryonic tissues were kept). Several embryos from the same litters were pooled by stage (10 embryos for E8.5, 6 embryos for E9.5, 5 embryos for E10.0 and 2 embryos for E10.5) (**Figure S1B**). To gain spatial resolution, embryos at E10.0 and E10.5 were divided into two parts along the A-P axis. Pooled samples were incubated on FACSmax Cell Dissociation Solution (Amsbio, T200100) with 10% Papain (30 U/mg, Sigma-Aldrich, 10108014001) for 11-12 min at 37°C to dissociate the cells as previously described ^61^. Samples were then resuspended in HBSS with 5% FBS, 10 µM rock inhibitor (Stemcell Technologies, Y-27632), and 1x non-essential aminoacids (Life Technologies, 11140050). Cells were filtered through 35 µm cell strainers (Falcon, 352235) and 20 µm filters (Miltenyi Biotec, 130-101-812). Samples with a cell viability above 60% were used for sequencing.

### 10X Genomics single-cell RNA-sequencing and quantification

Single-cell suspensions were loaded into the 10X Genomics Chromium Chips and followed the manufacturer’s instructions for Chromium Next GEM Single Cell 3’ Kit v3.1 (PN-1000121). Libraries were prepared using the Single Index Kit T Set A (PN-1000213) and sequenced on the Illumina HiSeq4000 system or the Illumina NovaSeq6000 system.

Gene expression matrices were computed from FastQ files using the kallisto (0.46.2) bustools (0.40.0) workflow ^102,103^. A custom reference genome was created using the Ensembl ASM229v1 genome which was supplemented with a pseudo-Y chromosome using known *M. domestica* cDNAs from Y-chromosome genes ^29,104^. Similarly, Ensembl gene models from release-103 were supplemented with predicted gene models for the Y chromosome. Sequencing replicates were merged before quantification. For quantification by kb count, the ‘10xv3’ technology was specified and data output in Cell Ranger format using the ‘cellranger’ option.

Cells were identified from droplets by DropletUtils (1.12.1) ^105^ using the equivalent threshold applied by Cell Ranger. Datasets were written in Cell Ranger format using the write10xCounts function.

### Seurat analysis

Datasets were loaded into Seurat (4.0.3) ^106^ objects for analysis in R (4.1.1). Initial quality control was applied to each sample independently using the number of genes identified, total UMI and percentage mitochondrial expression. The thresholds applied are listed in **Table S1**. A total of 23,669 cells were retained after filtering.

Cell cycle was assessed in each cell using the CellCycleScoring function and homologues to the S and G2M genes included in the Seurat cc.genes list. The difference (S-G2M) was calculated and saved into the object’s metadata. Similarly, the AddModuleScore function was used to identify male cells based on the expression of the Y-linked genes using *RPL10Y* and *HMGB3Y*.

Datasets were normalised and scaled using NormalizeData and ScaleData for visualisaiton and calculation of module and cell cycle scores; and SCTransform, for datasets that would be integrated. The 3,000 most-variable genes selected by FindVariableFeatures using the ‘vst’ method were used by SCTransform. Both the cell cycle score and Y-gene score were regressed from the data using the ‘vars.to.regress’ parameter of SCTransform.

Anterior and posterior datasets were integrated within time points using the Seurat IntegrateData workflow and the SCTransform-normalised data.

PCA was applied to each of the samples and integrated datasets and the first 70 principle components retained. Dimensionality was estimated for each dataset and is detailed in **Table S2**. UMAP and tSNE reductions were subsequently calculated using RunUMAP and RunTSNE, respectively. Clusters were identified using the FindNeighbors and FindClusters functions at a range of resolutions.

Datasets were subsequently subset to select embryonic cells, removing extraembyonic cells from the data (except in the E8.5 dataset). Once subset, the variable features were identified, the data rescaled, PCA recalculated and dimensionality estimated. UMAP and tSNE reductions and clusters could then be identified in each cell subset.

### Cluster annotation

Clusters were identified using gene module scores based on the combinatorial expression of conserved genes associated with specific cell states (**Table S3**). Module scores for cell types were calculated using AddModuleScore to help visualise cell types and select an appropriate cluster resolution for each dataset.

The early mesoderm clusters from the E10.0 dataset expressed mesoderm-associated markers including *TBXT, SNAI1*/*2, HAND1*/*2* and *MESP1* ^37,107–109^ but they did not have a clear signature of more mature subtypes. We took advantage of trajectory inference analyses using slingshot ^110^ to order these clusters and annotate them as nascent, early and advanced mesoderm (**Figure S4B**).

### Cross-species transcriptome-wide comparisons

Mouse ^17,19^, macaque ^25^ and rabbit ^21^ scRNA-seq datasets were downloaded and prepared for comparison to the opossum data. Each dataset was provided with cell type annotations which were kept for this analysis. Seurat objects were created for each dataset and only 1:1 orthologous genes, identified using the biomaRt package ^111^, were retained. For each species and time point, a set of variable genes were identified. For each time point comparison, the intersect between opossum and the comparison species’ variable genes were used to calculate Spearman’s rank correlation coefficient implemented by cor.test in R.

### A-P priority score

A-P priority score is calculated based on the difference between the number of differentially expressed genes (DEG) between the anterior dataset of time point t1 vs timepoint t0 and the number of DEG between the posterior dataset of time point t1 vs time point t_0_ and divided by the number of DEG between whole embryos of time points t_1_ vs t_0_.

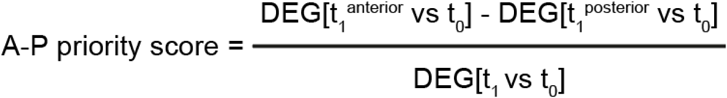

The number of DEG was calculated using the FindMarkers function (logfc.threshold = 0, min.pct = 0.005) from Seurat ^106^.

### HCR RNA-FISH

Fluorescent *in situ* hybridization chain reaction (HCR RNA-FISH) was performed using commercial reagents (Molecular Instruments) following the v3 protocol for whole-mount mouse embryos with some modifications ^112^. Briefly, embryos were fixed in 4% paraformaldehyde (PFA) for 2 hours at room temperature, dehydrated in a methanol series and stored at -20 °C until used. After rehydration, embryos were treated with proteinase-K (10 µg/mL) for 10 min at room temperature and post-fixed with 4% PFA for 20 min at room temperature. Embryos were then incubated with probe hybridization buffer for 30 min at 37 °C. Probes, designed by Molecular Instruments, were prepared at 4 pmol/mL in hybridization buffer. Embryos were incubated in the probe solution at 37 °C overnight. Excess probes were removed by washing embryos in probe wash buffer at 37 °C followed by washes in SSCT (5X sodium chloride sodium citrate, 0.1% Tween-20) at room temperature. Hairpins were prepared by snap-cooling at 95 °C for 90 s and cooled to room temperature protected from light and diluted in amplification buffer (60 pmol/mL). After 5 min incubation in amplification buffer, embryos were incubated in hairpin solution overnight at room temperature. Excess hairpins were removed by washing in SSCT. Nuclei were stained by DAPI and samples were mounted on slides using vectashield (Vector laboratories, H-1000-10). At least two opossum embryos were used per experiment.

### Cryosectioning

After whole-mount HCR RNA-FISH, selected opossum embryos were embedded in OCT and snap frozen. Transversal sections (20 µm thickness) were taken along the A-P axis and put onto Superfrost Plus slides (Thermo Scientific).

### Imaging

Whole-mount and sectioned embryos were imaged using a 40x immersion lens on an Olympus SoRa Spinning disc. Z-stacks were collected at 3-4 µm intervals, and tile scanning was used. Images were processed using Fiji ^113^. Maximum intensity Z-projections are shown.

## Acknowledgments

We thank the Francis Crick Institute Biological Research Facility, Advanced Sequencing, Scientific Computing, Advanced Light Microscopy, Experimental Histopathology, and Research Illustration for their assistance. We thank Güneş Taylor for advice on HCR RNA-FISH; Teresa Rayon, Malcolm Logan, James Briscoe, and members of the J.M.A.T. laboratory for discussion and feedback. Work in the J.M.A.T. and K.K.N. laboratories is supported by the Francis Crick Institute which receives its core funding from Cancer Research UK (CC2052, FC001120), the UK Medical Research Council (CC2052, FC00120), and the Wellcome Trust (CC2052, FC00120). Work in the J.M.A.T. and K.K.N. laboratories is also supported by the Wellcome Trust (222535/Z/21/Z and 221856/Z/20/Z). S.M. was supported by the Human Frontier Science Program (LT000293/2020-L2). For the purpose of Open Access, the authors have applied a CC BY public copyright licence to any Author Accepted Manuscript version arising from this submission.

## Author contributions

S.M. and J.M.A.T. conceived and designed the project. S.M. performed embryo experiments and bioinformatic analysis. C.B. performed bioinformatic analysis. G.A.L. performed initial quality control on datasets. W.V. advised on bioinformatic analysis. K.K.N. advised on data interpretation. S.M. and J.M.A.T. wrote the manuscript with input from all authors.

## Declaration of interests

The authors declare no competing interests.

## Notes

### Competing Interest Statement

The authors have declared no competing interest.

