## Supplemental Figures for "Marsupial single-cell transcriptomics provides an atlas of developmental heterochrony"

### Supplemental information

**Figure S1.** Characterisation of the opossum single-cell RNA-sequencing dataset.
**Figure S2.** Cross-species transcriptional correspondence.
**Figure S3.** Cluster annotation based on module scores in opossum E8.5 and E9.5 datasets.
**Figure S4.** Cluster annotation based on module scores in opossum E10.0 dataset.
**Figure S5.** Cluster annotation based on module scores in opossum E10.5 dataset.
**Figure S6.** Neural crest cells in the opossum embryo.
**Figure S7.** Spinal cord and neurons in the opossum embryo.
**Figure S8.** Patterning of the opossum forelimb.
**Figure S9.** The opossum gut.

A

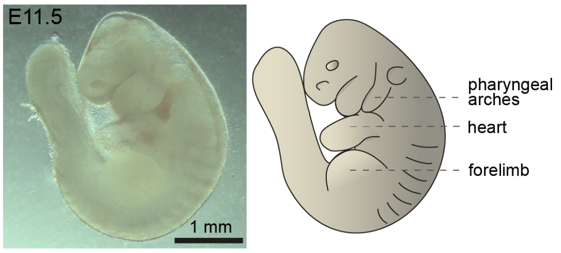


B

| dataset | number of embryos | number of cells pre QC | number of cells post QC | average number of genes | average number of UMI |
|---|---|---|---|---|---|
| E8.5 | 10 | 1,182 | 1,160 | 1,985 | 11,471 |
| E9.5 | 6 | 6,395 | 5,872 | 2,233 | 11,719 |
| E10.0 anterior | 5 | 4,249 | 3,994 | 1,655 | 7,361 |
| E10.0 posterior | 5 | 3,312 | 3,248 | 2,235 | 9,859 |
| E10.5 anterior | 2 | 4,373 | 4,218 | 2,837 | 15,028 |
| E10.5 posterior | 2 | 5,443 | 5,177 | 2,173 | 12,119 |
| | total | 24,954 | 23,669 | 2,186 | 11,260 |

C

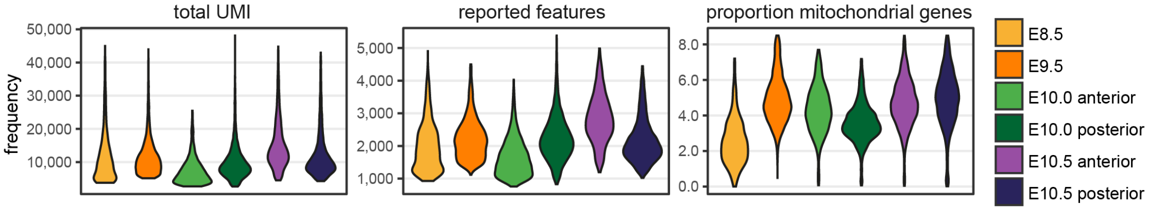


D

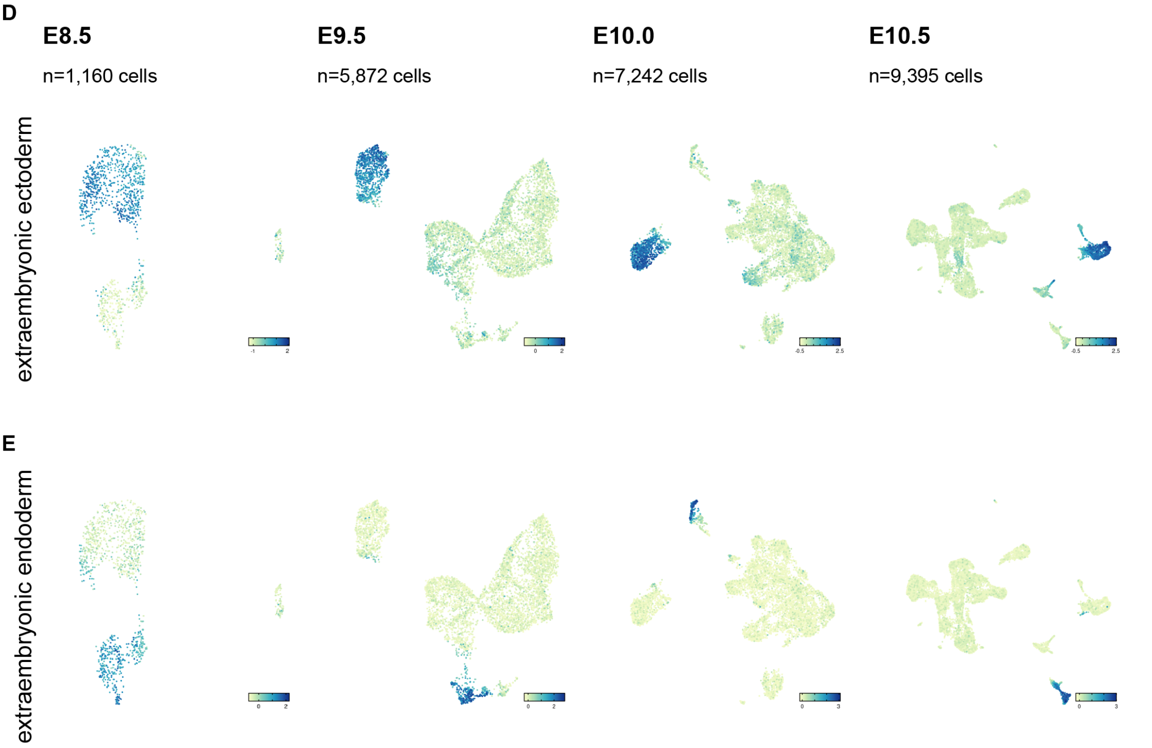


**Figure S1. Characterisation of the opossum single-cell RNA-sequencing dataset.**

(A) Stereo microscope image (left) and schematic illustration (right) of an E11.5 opossum embryo. Scale bar, 1 mm.

(B) Table with details about each scRNA-seq dataset including number of embryos, number of cells pre and post quality control (QC), average number of genes detected and average number of UMI (Unique Molecular Identifiers) detected.

(C) Distribution of number of UMI, number of features and percentage of mitochondrial genes in each single-cell dataset after QC filtering.

(D-E) Feature plots of opossum UMAP by time point showing expression of module scores associated with extraembryonic ectoderm (D) and extraembryonic endoderm (E). Module scores are based on the combinatorial expression of markers reported in Table S3.

Figure S2

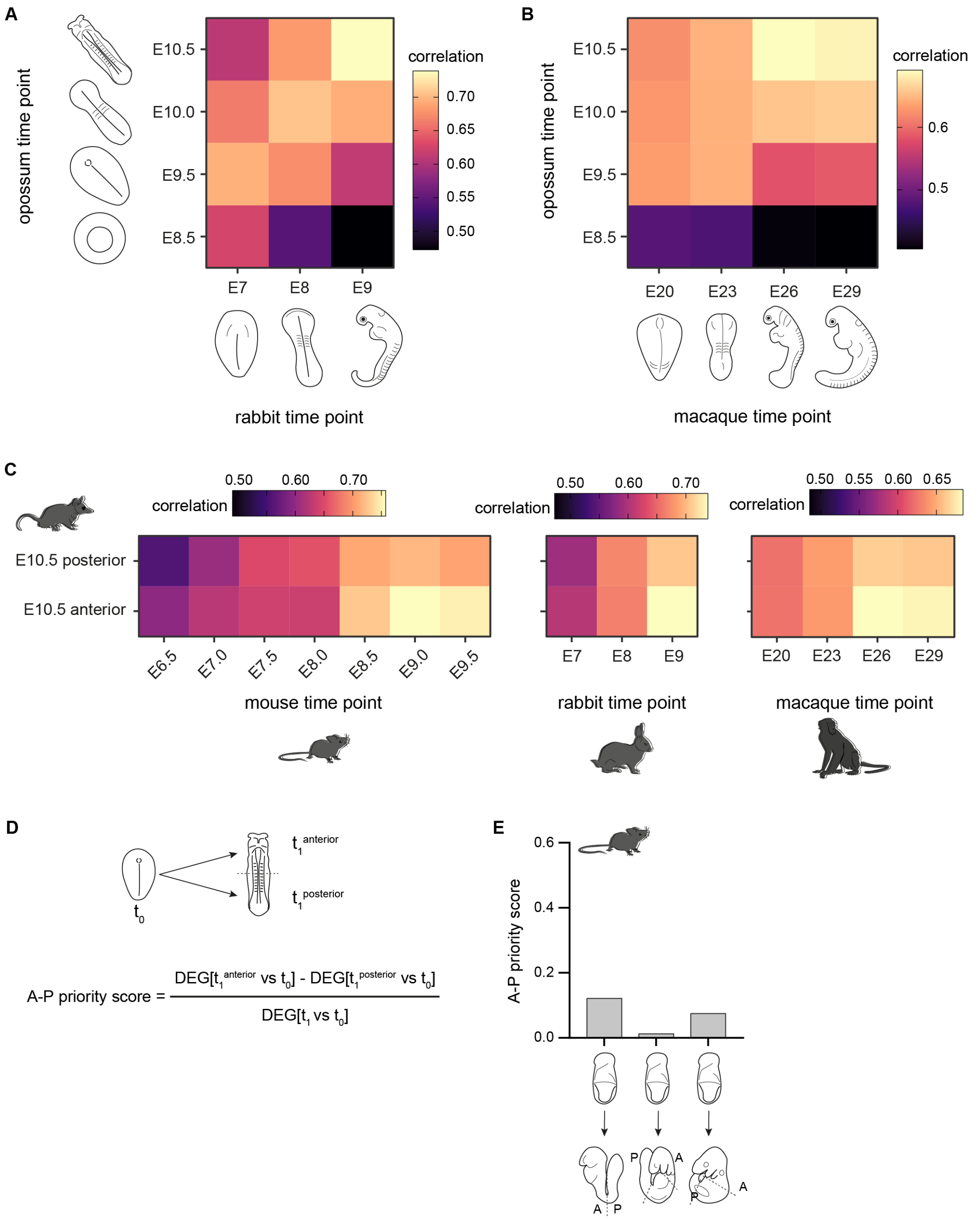


**Figure S2. Cross-species transcriptional correspondence.**

(A) Heatmap of Spearman's rank correlation coefficients of gene expression per opossum (vertical) and rabbit (horizontal) time point.

(B) Heatmap of Spearman's rank correlation coefficients of gene expression per opossum (vertical) and macaque (horizontal) time point.

(C) Heatmap of Spearman's rank correlation coefficients of gene expression per opossum E10.5 anterior and posterior datasets versus mouse (left panel), rabbit (middle panel) and macaque (right panel) datasets by time point. The intersect of variable features across timepoints between both species is used to calculate the correlation in all comparisons (A, B and C).

(D) A-P priority score diagram. A-P priority score is calculated based on the number of differentially expressed genes (DEG) between the anterior dataset of time point $t_1$ vs timepoint $t_0$ minus the number of DEG between the posterior dataset of time point $t_1$ vs time point $t_0$ and divided by the number of DEG between whole embryos of time points $t_1$ vs $t_0$.

(E) A-P priority score for mouse embryos using E7.5 as $t_0$ and different end time points ($t_1$ = E8.5, E9.0 and E9.5).

**A E8.5**

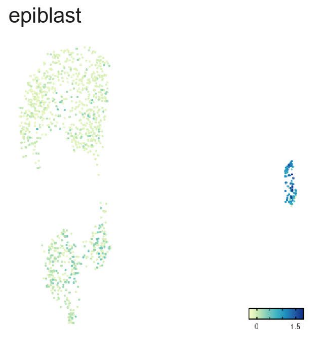


**B E9.5**

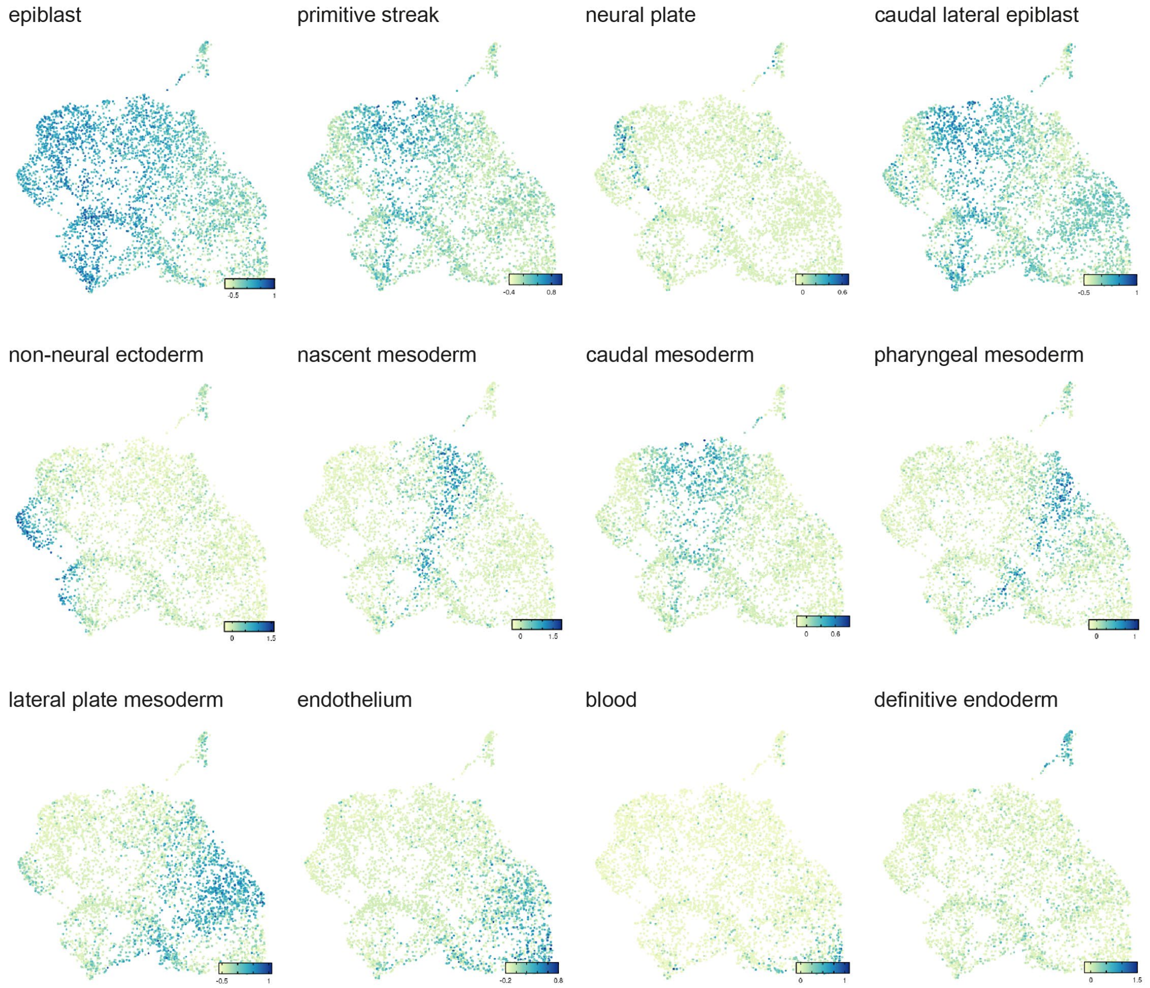


**Figure S3. Cluster annotation based on module scores in opossum E8.5 and E9.5 datasets.**

(A) Feature plot of opossum E8.5 UMAP showing expression of module score associated with epiblast cells.

(B) Feature plots of opossum E9.5 UMAP showing expression of module scores associated with selected cell types. Module scores are based on the combinatorial expression of markers reported in Table S3.

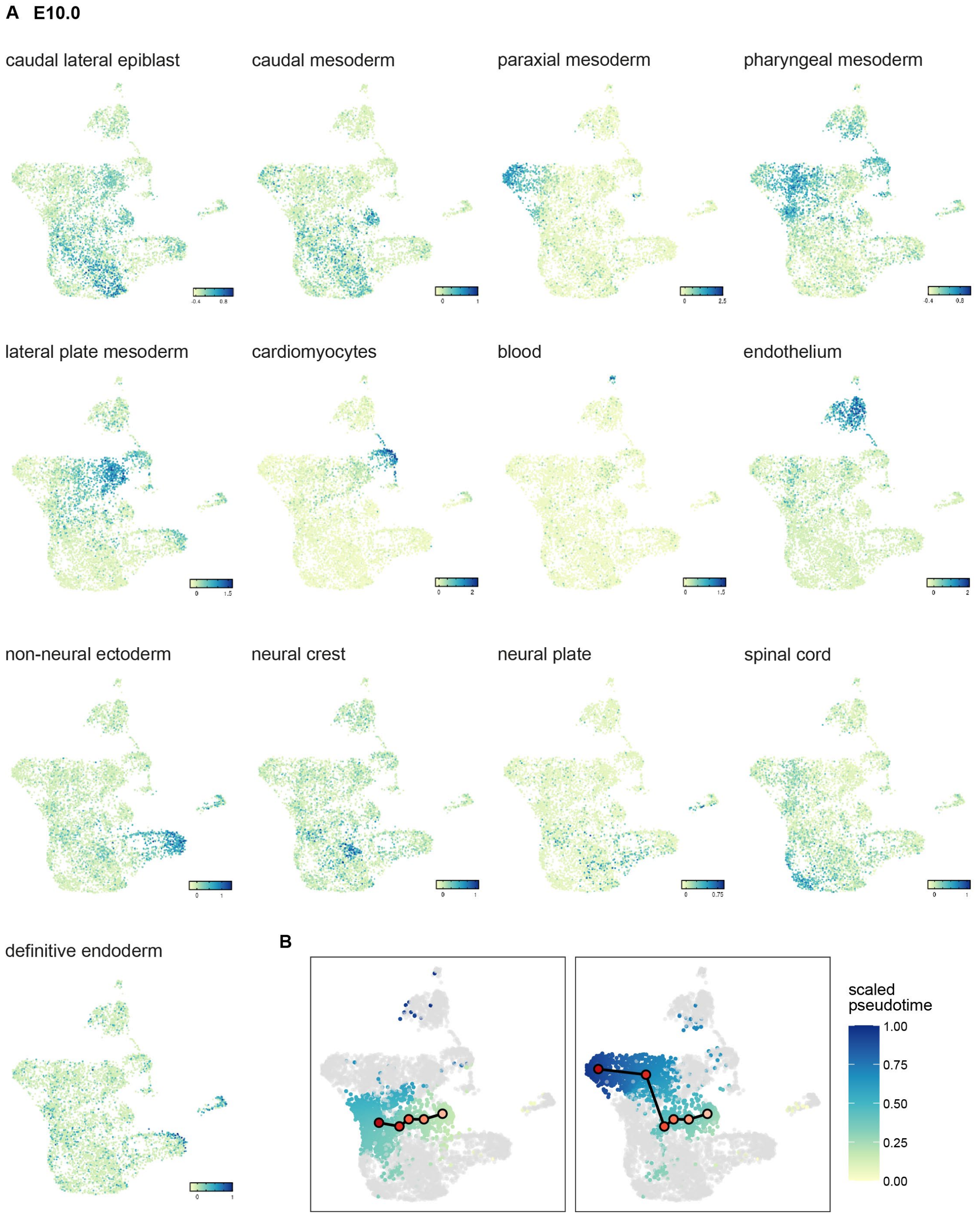


**Figure S4. Cluster annotation based on module scores in opossum E10.0 dataset.**

(A) Feature plot of opossum E10.0 UMAP showing expression of module score associated with selected cell types. Module scores are based on the combinatorial expression of markers reported in Table S3.

(B) Opossum E10.0 UMAP showing pseudotime of early mesoderm clusters. Cells from selected clusters are coloured in a gradient from yellow to dark blue to illustrate pseudotime. The average pseudotime of each cluster is also represented with a circle coloured from yellow (early) to red (late).

**A E10.5**

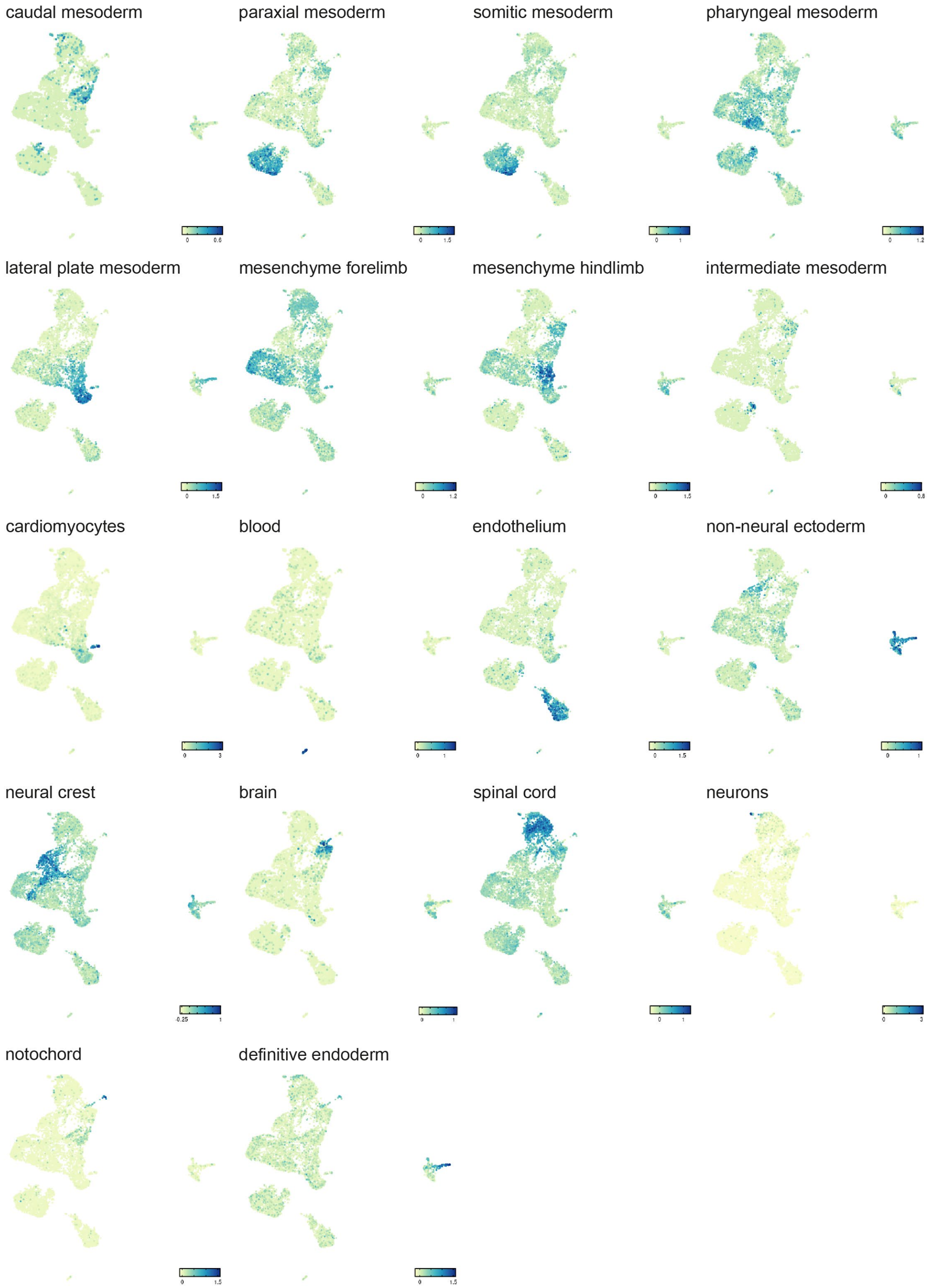


**Figure S5. Cluster annotation based on module scores in opossum E10.5 dataset.**

(A) Feature plot of opossum E10.5 UMAP showing expression of module score associated with selected cell types. Module scores are based on the combinatorial expression of markers reported in Table S3.

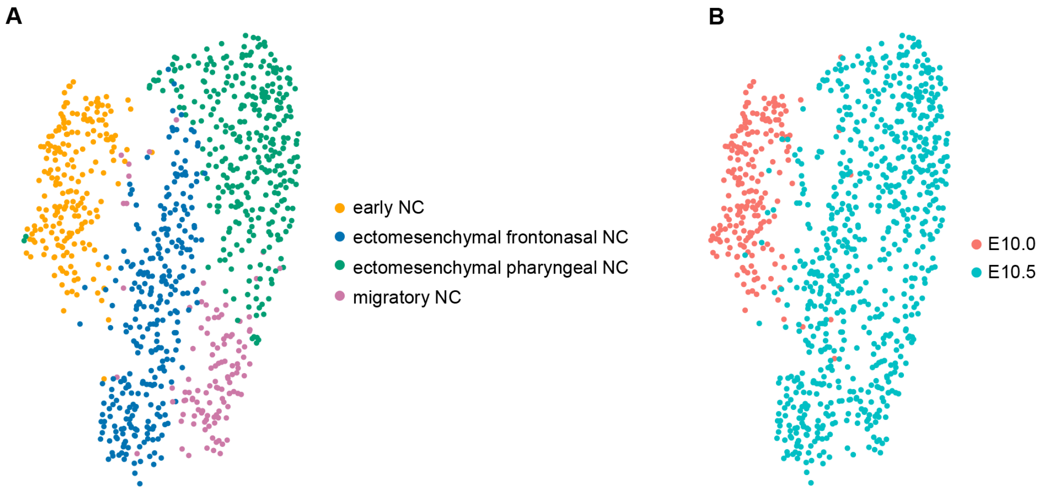


**Figure S6. Neural crest cells in the opossum embryo.**
(A) UMAP of anterior neural crest cells from E10.0 and E10.5 embryos. Cluster annotation based on the expression of key marker genes as shown in Figure 3A.
(B) UMAP of anterior neural crest cells coloured by time point.

Figure S7

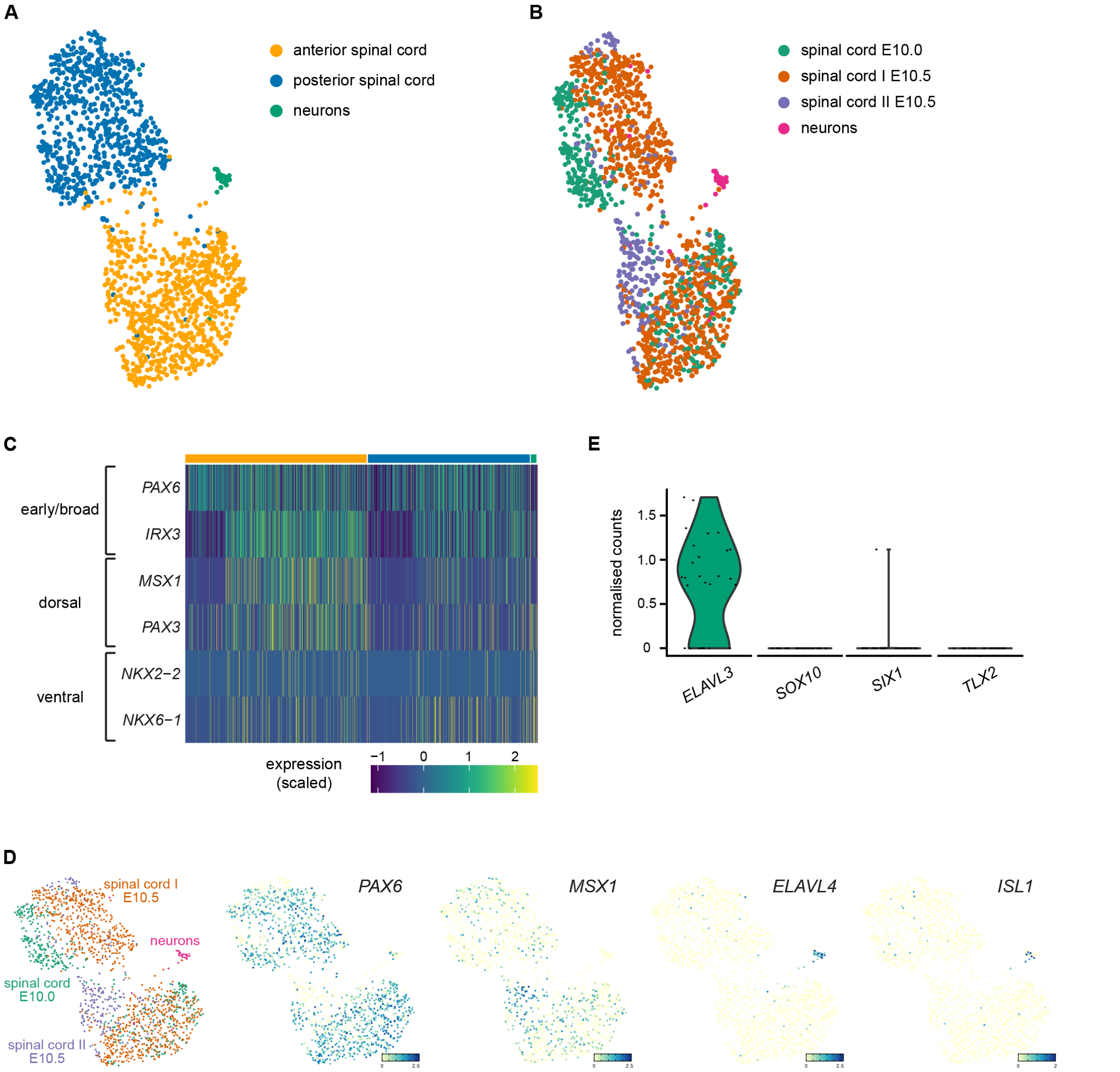


**Figure S7. Spinal cord and neurons in the opossum embryo.**

(A) UMAP of spinal cord cells and neurons from E10.0 and E10.5 embryos. Cluster annotation based on the expression of *HOX* genes as shown in Figure 4A.

(B) UMAP of spinal cord cells and neurons coloured by cluster of origin in E10.0 and E10.5 datasets.

(C) Heatmap of expression of selected genes associated with early/broad (*PAX6*, *IRX3*), dorsal (*MSX1*, *PAX3*) and ventral (*NKX2-2*, *NKX6-1*) populations in spinal cord clusters (yellow = anterior spinal cord; blue = posterior spinal cord; green = neurons).

(D) Feature plots of spinal cord and neurons UMAP showing expression of key genes associated with early spinal cord (*PAX6*), dorsal spinal cord (*MSX1*), pan neurons (*ELAVL4*) and motor neurons (*ISL1*).

(E) Violin plots showing expression of pan-neuronal (*ELAVL3*) and neural crest derived neuronal markers (*SOX10*, *SIX1*, *TLX2*) in opossum neurons at E10.5.

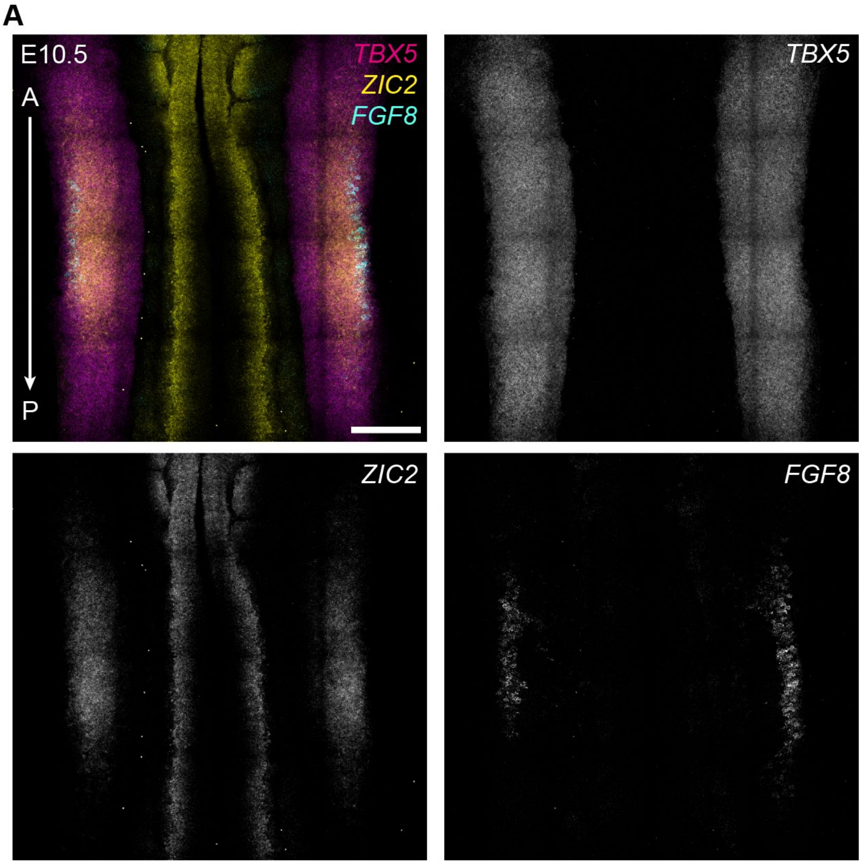


**Figure S8. Patterning of the opossum forelimb.**

(A) Expression of *TBX5*, *ZIC2* and *FGF8* by HCR RNA-FISH in the forelimb domain of an opossum E10.5 embryo. *ZIC2* marks the distal forelimb domain while *FGF8* is expressed in the limb ectoderm. Image is a maximal projection. Scale bar, 200 μm. A = anterior, P = posterior.

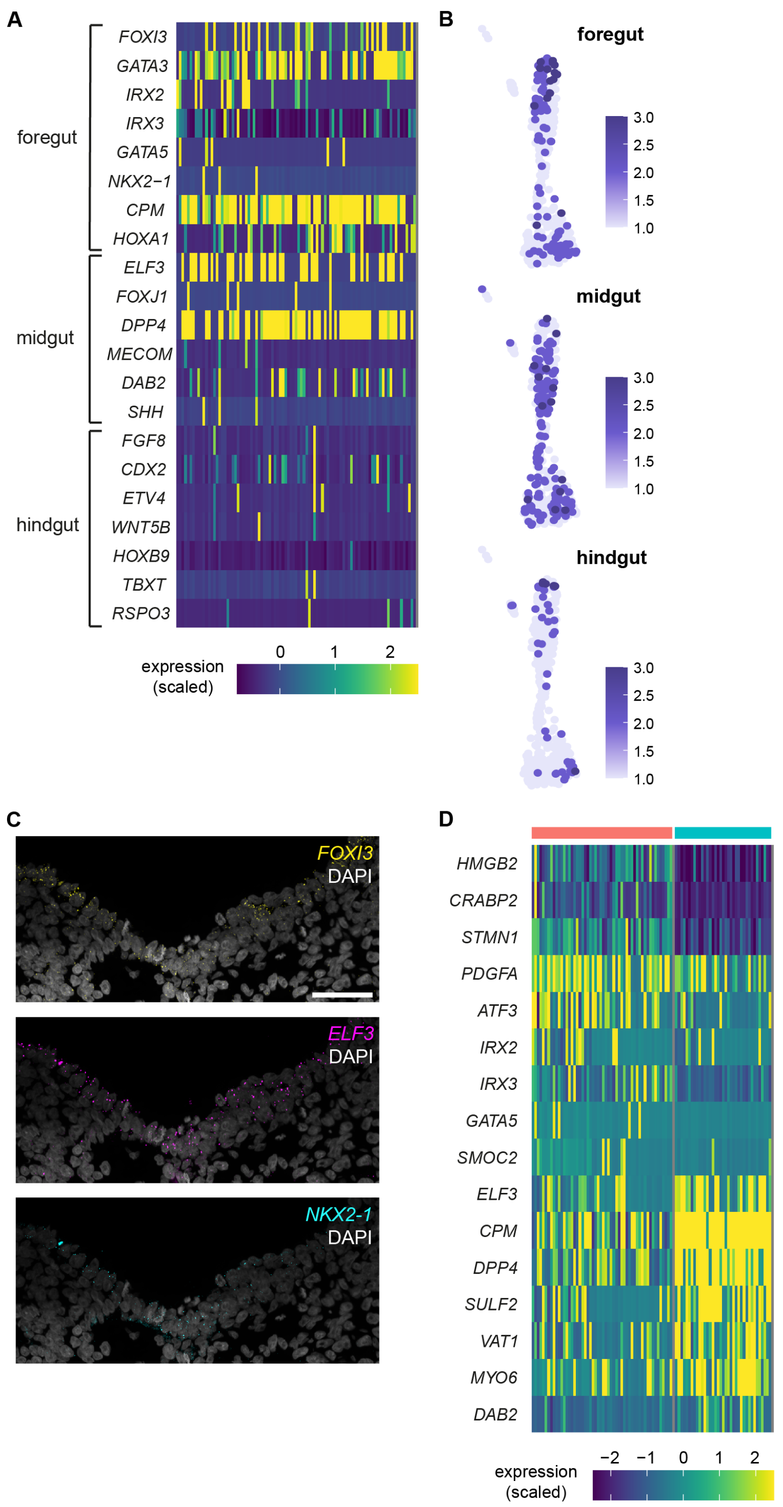


**Figure S9. The opossum gut.**

(A) Heatmap of expression of genes associated with foregut, midgut and hindgut identity in the definitive endoderm cells of E10.5 embryos.

(B) Feature plots of E10.5 extraembryonic endoderm cells showing enrichment of module scores associated with foregut, midgut and hindgut signatures. Module scores are based on the combinatorial expression of markers reported in Table S5.

(C) Expression of *FOXI3*, *ELF3* and *NKX2-1* by HCR RNA-FISH in the ventral foregut endoderm of an opossum E10.5 embryo. Image is a maximal projection. Scale bar, 50 μm.

(D) Heatmap of expression of genes enriched in different clusters of the E10.5 definitive endoderm cells. Cluster 1 (red bar) is enriched in genes associated with thymus, thyroid and lung progenitors while cluster 2 (blue bar) is enriched in genes associated with pancreas and small intestine progenitors.
